# Assessment of genetic diversity and SNP marker development within peanut germplasm in Taiwan by RAD-seq

**DOI:** 10.1101/2022.03.12.484056

**Authors:** Yu-Ming Hsu, Sheng-Shan Wang, Yu-Chien Tseng, Shin-Ruei Lee, Hsiang Fang, Wei-Chia Hung, Hsin-I Kuo, Hung-Yu Dai

## Abstract

The cultivated peanut (*Arachis hypogaea* L.) is an important oil crop but has a narrow genetic diversity. Molecular markers can be used to probe the genetic diversity of various germplasm. In this study, the restriction site associated DNA (RAD) approach was utilized to sequence 31 accessions of Taiwanese peanut germplasm, leading to the identification of a total of 17,610 single nucleotide polymorphisms (SNPs). The origin of these 31 accessions is contrasted so the global subset (n = 17) has greater genetic diversity than the local subset (n = 14). Concerning botanical varieties, the var. *fastigiata* subset has greater genetic diversity than the other two subsets of var. *vulgaris* and var. *hypogaea*, suggesting that novel genetic resources should be introduced into breeding programs to enhance genetic diversity. Principal component analysis (PCA) using genotyping data separated the 31 accessions into three clusters largely according to the botanical varieties, consistent with the PCA result for 282 accessions genotyped by 14 kompetitive allele-specific PCR (KASP) markers developed in this study. The SNP markers identified in this work not only revealed the genetic relationship and population structure of current germplasm in Taiwan, but also offer an efficient tool for breeding and further genetic applications.

## Introduction

Originated from South America, the cultivated peanut (*Arachis hypogaea* L.) is an allotetraploid (AABB, 2n=4x=40) and an important legume crop worldwide. Humans benefit from peanut seeds as food and source of oil due to their high percentage of proteins and fatty acids^1^. The annual production of peanuts has increased in the past 20 years to reach 53 million tons in 2020 according to FAOSTAT (http://www.fao.org/faostat). To fulfill the increasing need of peanut demand under the threat of climate change, breeding new varieties is an effective strategy to improve peanut qualitative and quantitative traits.

The conservation of *Arachis* germplasm and exploitation of their genetic diversity are crucial for the breeding program of the cultivated peanut. Presently, several gene banks are famous for their *Arachis* germplasm including the International Crops Research Institute for the Semi-Arid Tropics (ICRISAT), United States Department of Agriculture (USDA), and the Oil Crops Research Institute of the Chinese Academy of Agricultural Sciences (OCRI-CAAS). More than 15,000, 9,000 and 8,000 accessions were collected in ICRISAT, USDA, and OCRI-CAAS^2^, respectively. On the other hand, understanding the genetic diversity of in-hand germplasm is the prerequisite before launching breeding programs, and the utilization of molecular markers is the predominant strategy to evaluate the genetic diversity of germplasm at present^3^. Cultivated peanut is known for its low genetic diversity due to the recent hybridization of its two ancestors and selection in breeding programs^4,5,6,7^. Even though the narrow genetic diversity of cultivated peanuts has hindered the development of molecular markers, simple sequence repeat (SSR) markers can be developed and utilized to assess the genetic diversity in cultivated peanut^8,9,10,11^. In particular, the population structures of 92 accessions in the US Peanut Mini Core Collection and 196 major peanut cultivars in China were revealed by SSR markers^12,13^. Although SSR markers were widely used for identifying genetic diversity of peanut populations, these studies had limited population size due to the challenging genotyping process.

Recently, the peanut genome projects made possible by next generation sequencing (NGS) have revolutionized genetic research in cultivated peanuts, so that the genomes of *Arachis hypogaea* L. and its two diploid ancestors, *A. duranensis* (AA) and *A. ipaensis* (BB), are now sequenced^6,7,14^. These high quality genome sequences have paved the way for developing high-throughput single nucleotide polymorphism (SNP) markers e.g. *via* genotyping-by-sequencing (GBS) that can then facilitate peanut molecular breeding. The 58K SNP array ‘*Axiom_Arachis*’, developed by resequencing 41 peanut accessions, was used to identify genetic diversity across 384 *Arachis* genotypes including USDA Mini Core Collection and wild species^15,16^, while 787 accessions from the U.S. Peanut core collection were genotyped by the 14K ‘*Arachis_Axiom2*’ SNP array to reveal their genetic diversity^17^. Compared with SNP arrays, GBS is a more cost-effective technique based on sequencing of the reduced genome associated with restriction sites by NGS^18,19^. In peanut research, this technique was applied in SNP development, enabling the construction of genetic maps for quantitative trait locus (QTL) mapping and the analysis of population structure^20,21,22^.

In Taiwan, peanut breeding programs can be traced back to the late 1950’s. To date, most varieties developed locally have been obtained by conventional breeding based on evaluating morphological traits. In such breeding programs, the parental selection mainly relied on the pedigree information or the source of collection to infer genetic relationships. Thus, exploiting available molecular tools to characterize the present peanut varieties in Taiwan should allow improved breeding programs in the future. In this study, we performed the restriction site-associated DNA (RAD) approach to sequence 31 genotypes - including current elite varieties developed in Taiwan and important accessions introduced from abroad - to reveal the underlying genetic diversity. Furthermore, 14 kompetitive allele-specific PCR (KASP) markers were designed and then used to genotype 282 other accessions. Overall, this work reveals the genetic structure of peanut germplasm in Taiwan through SNP markers identified by RAD-seq and these markers can be used for a number of applications such as variety identification and breeding programs.

## Materials and methods

### Plant materials and DNA extraction

31 accessions, maintained by Taiwan Agricultural Research Institute (TARI) and Tainan District Agricultural Research and Extension Station (Tainan DARES), were chosen for RAD-seq construction. These accessions consist of elite cultivars, advanced breeding lines, and “introduced” old accessions acquired in South American countries close to the origin of peanut (Supplementary Table S1). Among 31 accessions, there are 13 Spanish, 10 Valencia, 4 Virginia, and 4 Runner type accessions. For the genotyping via KASP markers, 282 accessions were obtained from the National Plant Genetic Resources Center in TARI, including 66 Spanish, 27 Valencia, 49 Virginia and 88 Runner type accessions. The DNA extraction of all accessions was based on the modified CTAB method using young leaves collected from seedlings within two weeks^23^.

### Phenotypic evaluation

24 out of the 31 accessions used in RAD-seq and 282 additional accessions of the germplasm were phenotyped in the fall of 2016 in TARI (coordinates 24°01’47.5”N 120°41’47.4”E), and 20 plants of each accession were evaluated for 8 quantitative traits, including days to flowering (between the sowing and flowering date), plant architecture, number of pods, yield, 100-pod weight, 100-seed weight, rust resistance and leaf spot resistance. The susceptibility to these two peanut diseases was quantified under natural conditions in the field since these diseases develop spontaneously during the fall, and the disease symptoms were scored from 1 (having no symptoms) to 9 (highly susceptible)^24^. Depending on the degree of inclination from verticality, plant architecture was scored from 0 (the most upright) to 9 (the most prostrate).

### RAD-seq library construction and SNP calling

After extracting DNA of 31 accessions based on the modified CTAB method, we used the QIAGEN kit (DNeasy Blood & Tissue Kit; Qiagen, https://www.qiagen.com/, Hilden, Germany) to perform the DNA purification and ensure DNA quality. These samples were then used to make two RAD-seq libraries from 16 and 15 accessions, respectively, following the published protocol^25^, using *PstI* as the digestion enzyme. Next-generation sequencing of each library was carried out in the Genome Research Center of Yang-Ming University using the Illumina Hiseq 2500 platform (Illumina Inc., https://www.illumina.com, CA, USA) with 100 bp single-end reads in two lanes. The sequencing data have been deposited at National Center for Biotechnology Information (NCBI) under BioProject PRJNA811600. For SNP calling, single-end reads were first debarcoded by Stacks using the program “process_radtags”^26^. Then, we used Burrows-Wheeler Alignment (BWA) v0.7.17-r1188 “aln” to align the reads of each accession onto the reference genome of cultivated peanut and its diploid ancestors for identifying SNPs used in the genetic diversity analysis and the development of KASP markers, respectively^6,14,27^. When the genome of cultivated peanut was published^14^, all of the KASP markers used in this study had already been designed using the merged genomes of two diploid ancestors of cultivated peanut^6^. Thus, identified SNPs based on the genome of cultivated peanut were only used for the *in-silico* analyses that investigated the genetic diversity of the 31 accessions, but in all cases the SNP calling pipeline was the same. After the alignment was finished, Samtools and BCFtools were utilized for SNP calling and filtering, SNPs were kept with (1) base quality ≥ 20, (2) mapping quality score ≥ 20, and (3) depth ≥ 3. Then, a customized R script was used to create Variant Call Format (VCF) files encompassing qualified SNPs that discriminated the 31 accessions^28^.

### The development and validation of KASP markers

Among SNPs available for distinguishing 31 accessions based on the genome of the two diploid ancestors^6^, we extracted 1,230 homozygous SNPs with informative alleles in all 31 accessions, and then discarded 783 SNPs having Polymorphic Information Content (PIC) values lower than the average over all SNPs. Finally, 29 out of 477 SNPs with an average PIC value of 0.28 were selected for developing KASP markers. These 29 putative SNPs with 100 bp flanking sequences on both sides were used for designing KASP primers that were then synthesized in LGC genomics (http://www.lgcgroup.com, Teddington, England). The validation of KASP markers was performed on the 96-well StepOnePlus™ Real-Time PCR System (Thermo Fisher Scientific Inc., https://www.thermofisher.com, DE, USA), and each 10-μL reaction consisted of 12.5 ng of DNA, 0.14 μL of KASP assay mix and 5 μL of KASP Master Mix (2X). The PCR protocol was carried out as follows: (1) pre-read stage at 30°C for 1 minute, (2) hold stage at 94°C for 15 minutes, (3) PCR stage 1 of 10 touchdown cycles using 94°C for 20 seconds and 61°C (decreasing 0.6°C per cycle) for 1 minute, (4) PCR stage 2 with 26 amplification cycles at 94°C for 20 seconds and 55°C for 1 minute, and (5) post-read stage at 30°C for 1 minute. When the PCRs were completed, the fluorescent signals of samples were analyzed by the StepOne™ software for determining genotypes.

### Statistical analysis

All statistical analyses were carried out in R 3.63. The PIC value and the expected heterozygosity (He) were determined for each SNP marker^29,30^. Principal component analysis (PCA) using phenotypic data was performed by the “PCA” function in the “FactoMineR” package^31^. In the “poppr” package, the “bitwise.dist” function was used to calculate the genetic distances between the 31 accessions, distances that depend on the fraction of loci which differ between germplasm^32,33^, and the “aboot” function was utilized to construct the dendrograms based on the unweighted pair group method with arithmetic mean (UPGMA) with 1000 bootstraps. In the “adegenet” package, PCA for SNP data from 31 accessions was carried out by the “glPCA” function, and population structure of 282 accessions was addressed by successive K-mean clustering and discriminant analysis of principal components (DAPC) using the “find.clusters” and “dapc” function, respectively^34,35^. In addition to the dendrogram plot, created by the “plot.phylo” function in the “ape” package^36^, all visualization was performed using the “ggplot” function in the “tidyverse” package^37^.

## Results

### SNP marker development from 31 peanut accessions using RAD-seq

In this study, 31 peanut accessions were chosen to conduct RAD-seq, of which 17 accessions were introduced from abroad and 14 accessions developed or collected in Taiwan. This collection has important agronomic traits including yield-related traits, resistances to biotic and abiotic stresses, and valuable characteristics considering the genetic diversity (Supplementary Table S1).

In the RAD-seq approach, the six-cutter enzyme, *PstI*, was utilized for the DNA digestion, and so sequencing of each accession focused on approximately 5 % (100-bp extensions on both side of a *PstI* cutting site that occurs every 4,096 bp on average) of the total cultivated peanut genome (2.7 Gb). The estimated sequencing depth in the 31 accessions ranged from 4.26 (HL2) to 15.01 (Red), and the average depth was 9.47. In addition, more than 99.0 % of sequenced reads from all samples were properly aligned to the reference genome. Compared to the reference genome, *A. hypogaea* cv. Tifrunner, there were 1,475 to 14,471 SNPs identified from these 31 accessions with an average of 5,249 SNPs, and more than 3 quarters of these polymorphisms are homozygous. In addition, the transition/transversion (Ts/Tv) ratio ranges from 0.48 to 1.19 (Table 1). In terms of the three botanical varieties of the cultivated peanut, accessions from subsp. *fastigiata* var. *vulgaris* (Spanish type), subsp. *fastigiata* var. *fastigiata* (Valencia type) and subsp. *hypogaea* var. *hypogaea* (Virginia/Runner types) have a total of 5,006, 5,119 and 5,905 SNPs, i.e., the differences across varieties are very small. Interestingly, while the 31 accessions were separated into the global and local collection corresponding to 17 genotypes introduced from other countries and 14 genotypes from Taiwan, respectively, the global collection has an average of 6,071 SNPs which is higher than the average of 4,252 SNPs for the local collection. Moreover, 8 introduced accessions, collected in South America close to the center of origin of cultivated peanut, lead to an average of 7,139 SNPs, even higher than that of the global collection, which suggests that the global collection germplasm from various countries have more polymorphisms than the local one containing mainly Taiwanese cultivars. Then, the next stage of filtration was performed to keep only SNPs differentiating these 31 accessions. As a result, 3,474 out of 17,610 SNPs were finally kept for the genetic diversity analysis using a tolerance of 6 missing values (20%) at most for each polymorphism.

**Table 1.** Sequence and SNP information of our 31 accessions in the Taiwanese peanut germplasm

| Germplasm | Mapped reads | The percentage of mapped reads | Estimated depth<sup>a</sup> | Filtered SNPs compared to *A. hypogaea* cv. Tifrunner | Transition/Transversion ratio | Homozygous SNPs compared to RF |
| --- | --- | --- | --- | --- | --- | --- |
| PI153169 | 10,886,987 | 99.44% | 8.4 | 5,666 | 0.66 | 5,342 |
| PI259717 | 11,647,977 | 99.26% | 8.98 | 5,502 | 0.67 | 5,118 |
| PI565455 | 13,808,309 | 98.99% | 10.65 | 14,471 | 0.63 | 14,051 |
| Tainung 7 (TNG7) | 15,599,384 | 99.52% | 12.04 | 6,166 | 0.6 | 5,848 |
| Tainung 10 (TNG10) | 15,727,918 | 99.56% | 12.14 | 5,884 | 0.64 | 5,630 |
| Tainan 14 (TN14) | 12,568,819 | 99.18% | 9.7 | 1,682 | 1.12 | 1,348 |
| Tainan 15 (TN15) | 8,267,427 | 99.11% | 6.38 | 1,475 | 1.19 | 1,217 |
| Tainan 18 (TN18) | 6,503,808 | 99.15% | 5.02 | 1,794 | 0.68 | 1,461 |
| Tainan Selection 9 (TNS 9) | 17,017,130 | 99.04% | 13.13 | 6,884 | 0.59 | 6,387 |
| Hualien 1 (HL1) | 12,662,231 | 99.14% | 9.77 | 3,600 | 0.67 | 3,230 |
| India | 13,828,557 | 99.09% | 10.67 | 3,495 | 0.72 | 3,174 |
| Xiamen | 14,721,052 | 99.02% | 11.35 | 5,858 | 0.7 | 5,483 |
| Vietnam | 10,331,877 | 99.10% | 7.97 | 2,605 | 0.75 | 2,301 |
| PI118480 | 11,987,640 | 99.48% | 9.25 | 5,564 | 0.83 | 5,277 |
| PI118989 | 14,285,571 | 99.28% | 11.02 | 11,823 | 0.53 | 11,367 |
| PI155112 | 10,196,688 | 99.44% | 7.87 | 5,163 | 0.74 | 4,829 |
| PI314817 | 11,876,221 | 99.38% | 9.16 | 11,051 | 0.73 | 10,653 |
| PI338337 | 13,887,142 | 99.62% | 10.72 | 6,187 | 0.69 | 5,830 |
| Tainan 16 (TN16) | 9,887,062 | 99.15% | 7.63 | 1,683 | 1.19 | 1,326 |
| Tainan 17 (TN17) | 6,187,833 | 99.19% | 4.77 | 1,606 | 1.14 | 1,324 |
| Hualien 2 (HL2) | 5,522,671 | 99.17% | 4.26 | 1,896 | 0.62 | 1,642 |
| E01001 | 8,193,785 | 99.22% | 6.32 | 2,376 | 0.98 | 1,923 |
| E01004 | 7,837,504 | 99.21% | 6.05 | 2,759 | 0.85 | 2,393 |
| Red | 19,447,996 | 99.17% | 15.01 | 6,204 | 0.51 | 5,797 |
| NS011001 | 12,917,315 | 99.15% | 9.97 | 4,147 | 0.87 | 3,735 |
| PI109839 | 13,211,606 | 99.57% | 10.19 | 6,805 | 0.48 | 6,499 |
| Taichung 1 (TC1) | 16,400,219 | 99.63% | 12.65 | 7,390 | 0.65 | 7,097 |
| PI145681 | 12,327,318 | 99.47% | 9.51 | 2,618 | 0.56 | 2,342 |
| PI599592 | 16,433,115 | 99.55% | 12.68 | 6,408 | 0.69 | 6,150 |
| PI203396 | 11,566,479 | 99.59% | 8.92 | 4,855 | 0.59 | 4,593 |
| Penghu 1 (PH1) | 14,732,914 | 99.34% | 11.37 | 9,115 | 0.58 | 8,713 |

<sup>a</sup>The estimated depth was calculated by the total number of bases divided by 4.8% of 2.7 Gb, the size of reduced reference genome.

### Evaluation of genetic diversity and cluster analysis based on 31 peanut accessions

The genetic diversity of the 31 peanut accessions was quantified by a number of measures, including the expected heterozygosity (He), the major allele frequency (MAF), polymorphic information content (PIC), and genetic distance. The genetic distance is based on bitwise distance, identical to Provesti’s distance, growing with the fraction of genetically different loci between 31 accessions^32^. The pairwise comparison of the genetic distance between accessions is listed in Supplementary table S2. On average, these 31 peanut accessions have a He of 0.19, PIC of 0.16, MAF of 0.86, and distance of 0.17. While considering botanical varieties, germplasm from subsp. *fastigiata* var. *fastigiata* have the largest average He, PIC and genetic distance (He = 0.18, PIC = 0.14, distance = 0.15) and smallest MAF (0.87), to be compared to that of the germplasm from subsp. *fastigiata* var. *vulgaris* (He = 0.13, PIC = 0.11, MAF = 0.90, distance = 0.11) or subsp. *hypogaea* var. *hypogaea* (He = 0.12, PIC = 0.10, MAF = 0.92, distance = 0.12) (Table 2), showing that Valencia-type germplasm acquired higher genetic diversity than both Spanish-type and Virginia/Runner-type germplasm. In terms of the collection source, the global collection has larger average He, PIC and genetic distance (He = 0.19, PIC = 0.15, distance = 0.17) and smaller MAF (0.86) than the local collection (He = 0.16, PIC = 0.13, MAF = 0.88, distance = 0.14), indicating that the global collection has greater genetic diversity than the local collection. In addition, the distance tree for cluster analysis was reconstructed based on the UPGMA method with 1,000 bootstraps (Fig. 1). The result showed that 26 out of the 31 accessions were clustered into 3 groups mainly according to three botanical varieties. Germplasm of subsp. *fastigiata* var. *fastigiata* and subsp. *fastigiata* var. *vulgaris* were clustered into Group I and II with a distance of 0.19, and germplasm of subsp. *hypogaea* var. *hypogaea* was clustered into Group III separated from Group I and II with a distance of 0.21. With the exception of 5 accessions, NS011001 (Virginia type) was clustered into group I with mainly Valencia type germplasm, while two Valencia type accessions, Red and HL2, were clustered into group II with mostly Spanish type germplasm. Interestingly, TN16 and TN17, two Valencia type cultivars, are clustered into group III, but they are separated from accessions of subsp. *hypogaea* var. *hypogaea* with a distance of 0.18.

**Table 2.** Genetic diversity in the 31 accessions of Taiwanese peanut germplasm

| group | number | mean He | mean MAF<sup>a</sup> | mean PIC<sup>b</sup> | mean genetic distance |
| --- | --- | --- | --- | --- | --- |
| The whole collection | 31 | 0.19 | 0.86 | 0.16 | 0.17 |
| The origin of germplasm |  |  |  |  |  |
| The global subset | 17 | 0.19 | 0.86 | 0.15 | 0.17 |
| The local subset | 14 | 0.16 | 0.88 | 0.13 | 0.14 |
| The botanical variety |  |  |  |  |  |
| var. *vulgaris* | 13 | 0.13 | 0.90 | 0.11 | 0.11 |
| var. *fastigiata* | 11 | 0.18 | 0.87 | 0.14 | 0.15 |
| var. *hypogaea* | 7 | 0.12 | 0.92 | 0.10 | 0.12 |

<sup>a</sup>MAF, major allele frequency
<sup>b</sup>PIC, polymorphic information content

**Figure 1.**
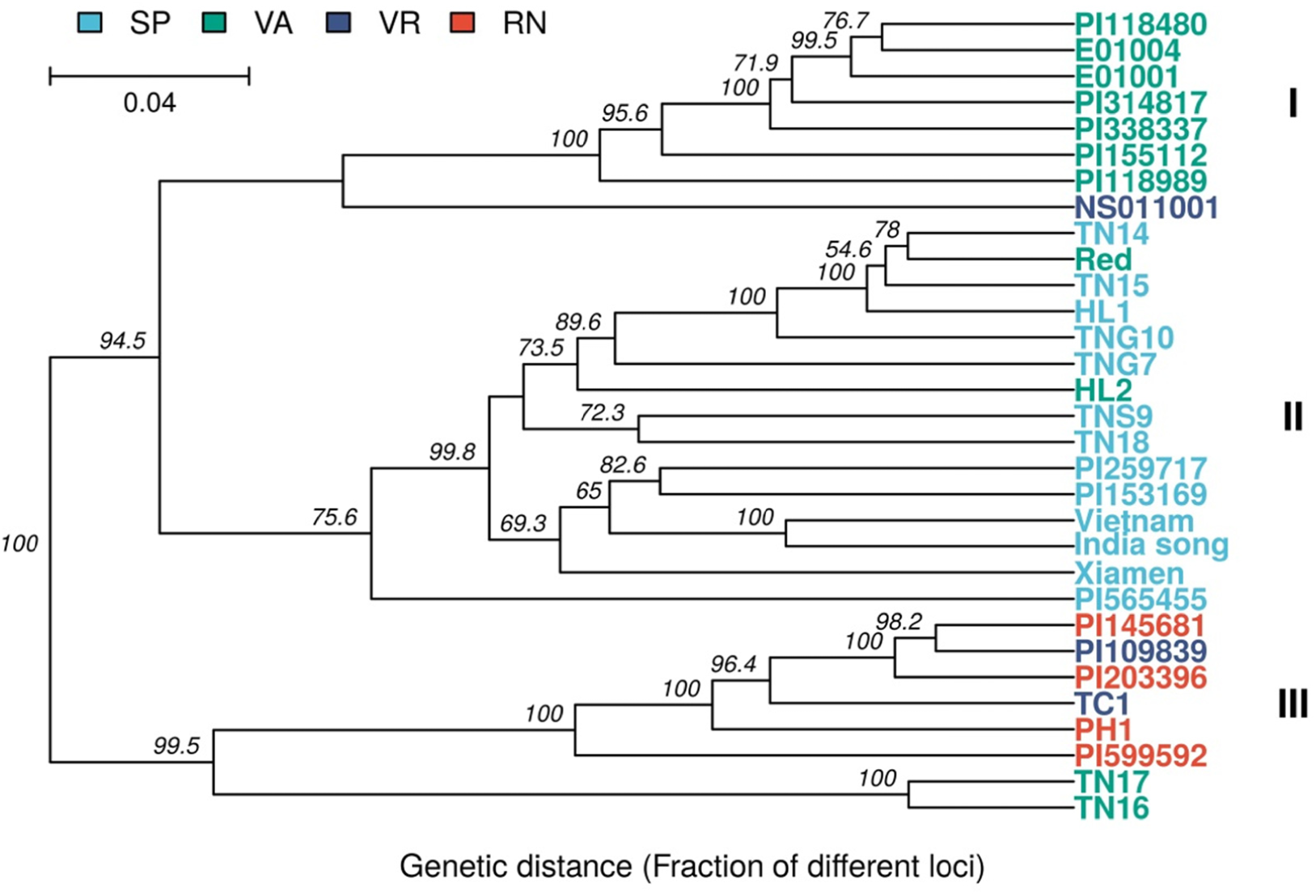
Dendrogram of the 31 accessions created from the unweighted pair group method with arithmetic mean (UPGMA). This dendrogram was based on the pairwise genetic distance with 1,000 replicates using 3,474 single nucleotide polymorphisms (SNPs). The branch length represents genetic distance, and the scale is on the top left of this figure. The numbers on the branches are bootstrap percentages. The legend sho s the color of four market types, Spanish (SP), Valencia (VA), Virginia (VR), Runner (RN), and three clades clustered in this plot were named as I, II and III.

To further investigate and compare the genetic relationship among these germplasm based on genomic and phenotypic data, PCA were separately performed using genetic distances between the 31 accessions calculated *via* 3,474 SNPs and using the phenotypic data for 8 agronomic traits from 24 of the 31 accessions based on field experiments conducted in the fall of 2016. For PCA based on distance obtained using SNPs, the result showed that the first three principal components (PC1, PC2 and PC3) explain 24.2%, 20.8% and 8.2% of the variance, respectively, totaling 53.2 % of the overall genetic distance variance (Fig 2). On the other hand, PC1, PC2 and PC3 in the PCA that relied on the phenotypic data of 8 agronomic traits explain 33.0%, 23.9% and 16.3% of the phenotypic variance, respectively, cumulating 73.2 % of the overall phenotypic variance (Supplementary Fig. S1). However, the scatter plots of PCs suggests that PCA based on genomic data can better distinguish the 31 accessions. In the two biplots of PC1/PC2 and PC1/PC3 based on PCA using 3,474 SNPs, the first pair can distinguish 31 accessions into three clear groups, mainly according to three botanical varieties, subsp. *fastigiata* var. *vulgaris* (Spanish type), subsp. *fastigiata* var. *fastigiata* (Valencia type) and subsp. *hypogaea* var. *hypogaea* (Virginia/Runner types), while the second pair is capable of separating TN16 and TN17 from three clusters assigned by the first pair (Fig. 2). On the contrary, for the two biplots from PCA depending on phenotypic data, neither PC1/PC2 nor PC1/PC3 can clearly provide evidence for structure within the 31 accessions (Supplementary Fig. S1).

**Figure 2.**
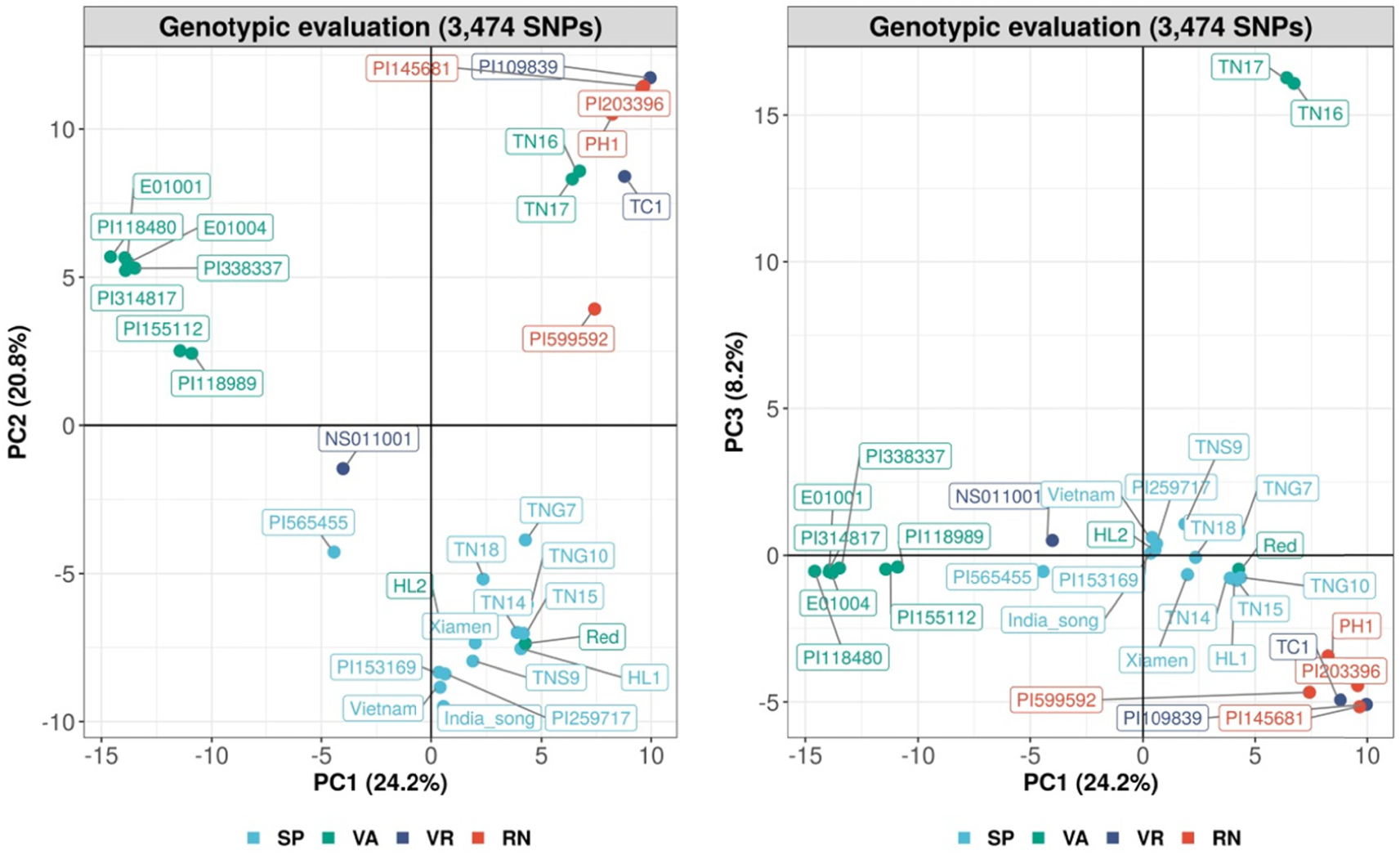
Principal component analysis (PCA) of the 31 accessions based on 3,474 single nucleotide polymorp**h**isms (SNPs). The 31 accessions were visualized by 4 market types, Spanish (SP), Valencia (VA), Virginia (VR), Runner (RN).

### The development and validation of KASP markers

In this study, one of our goals was to design a set of non-gel based SNP markers which can be exploited to investigate the genetic relationship between the germplasm collection conserved in the National Plant Genetic Resources Center of TARI. When this project was launched, the genome of cultivated peanut wasn’t published yet. Thus, the development of SNP markers for the KASP part of this work relied on the two diploid ancestors of cultivated peanut^6^. Note that the SNP calling pipeline used here was the same as the one that identified SNPs from the cultivated peanut genome for assessing the genetic diversity of the 31 accessions. Of the SNPs identified by the mapping to the two diploid ancestral genomes, 1,230 had both alleles represented in the 31 accessions while satisfying the constraint of being homozygous and having no missing data therein. 477 of these SNPs were kept because their PIC value was higher than the average one of the 1,230 homozygous SNPs (Supplementary File S1). 29 out of 477 SNPs with an average PIC value of 0.28 were selected for designing KASP markers. Based on the RAD-seq result, these 29 SNPs were able to group 31 accessions into three clusters according to their botanical varieties (Fig. 3). To validate these KASP markers, 282 accessions of the TARI germplasm with 66 Spanish, 27 Valencia, 49 Virginia and 88 Runner type accessions were genotyped, and 14 out of 29 KASP markers showed the stable and discernible genotyping result in the initial validation process.

**Figure 3.**
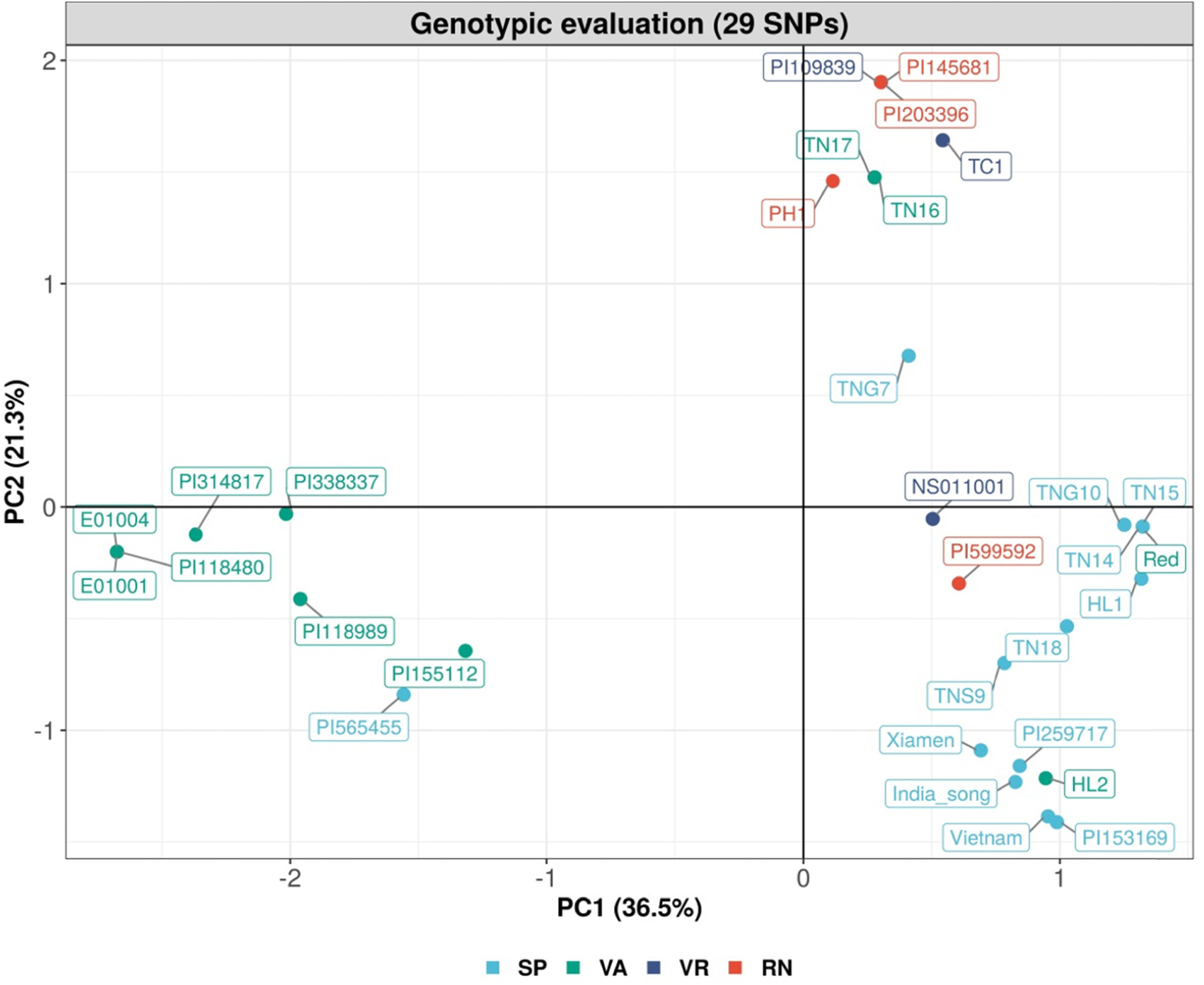
Principal component analysis (PCA) of the 31 accessions based on 29 single nucleotide polymorphisms (SNPs). 31 accessions were visualized by 4 market types, Spanish (SP), Valencia (VA), Virginia (VR), Runner (RN).

The population structure analysis of 282 accessions were then identified by PCA using genetic distances between these 282 accessions calculated by the KASP-marker genotyping data (Supplementary Table S3) and the phenotypic data from the field experiment in the fall of 2016 based on 8 agronomic traits. The two independent PCA biplots indicate that the one using the genotypic data performed better than the one using the phenotypic data to distinguish 282 accessions. For the scatter plot based on genomic data, PC1 and PC2 explain 37.1% and 18.4% of the variance of the genotyping data and can separate these accessions into 3 groups according to three botanical varieties (subsp. *fastigiata* var. *vulgaris*, subsp. *fastigiata* var. *fastigiata* and subsp. *hypogaea* var. *hypogaea*). On the other hand, the first two PCs from the PCA using phenotypic data account for 28.4% and 24.1% of phenotypic variation, and only roughly separate these accessions into two groups (subsp. *fastigiata* var. *vulgaris*/var. *fastigiata* and subsp. *hypogaea* var. *hypogaea*). Most Spanish and Valencia type accessions were difficult to distinguish using phenotypic data, and the grouping between Spanish/Valencia and Virginia/Runner accessions was less clear than the PCA result based on genotyping data (Fig. 4). To further identify the population structure within the 282 accessions, discriminant analysis of principal components (DAPC) was performed based on increasing number of clusters (K) assigned by successive K-means. In such an approach, the Bayesian information criterion (BIC) was used to assess the model relevance, and the result showed that it was best to go to ks of at least 3 for clustering the 282 accessions (Figure S2). Similarly, DAPC was conducted for 2, 3 and 4 clusters to explore the population structure of 282 accessions. At k=2, clusters correspond to Spanish/Valencia and Virginia/Runner type accessions. At k=3, the 3 groups correspond largely to Spanish, Valencia and Virginia/Runner. At k=4, the overall grouping trend is similar to that with k=3, but have a mixture of accessions from the three botanical varieties that are assigned into the fourth group (Fig. 5). This result suggests that our KASP markers are effective for identifying the population structure of peanut germplasm according to the botanical varieties, and it can even illustrate the similar genetic background acquired by accessions corresponding to a mixture of botanical varieties.

**Figure 4.**
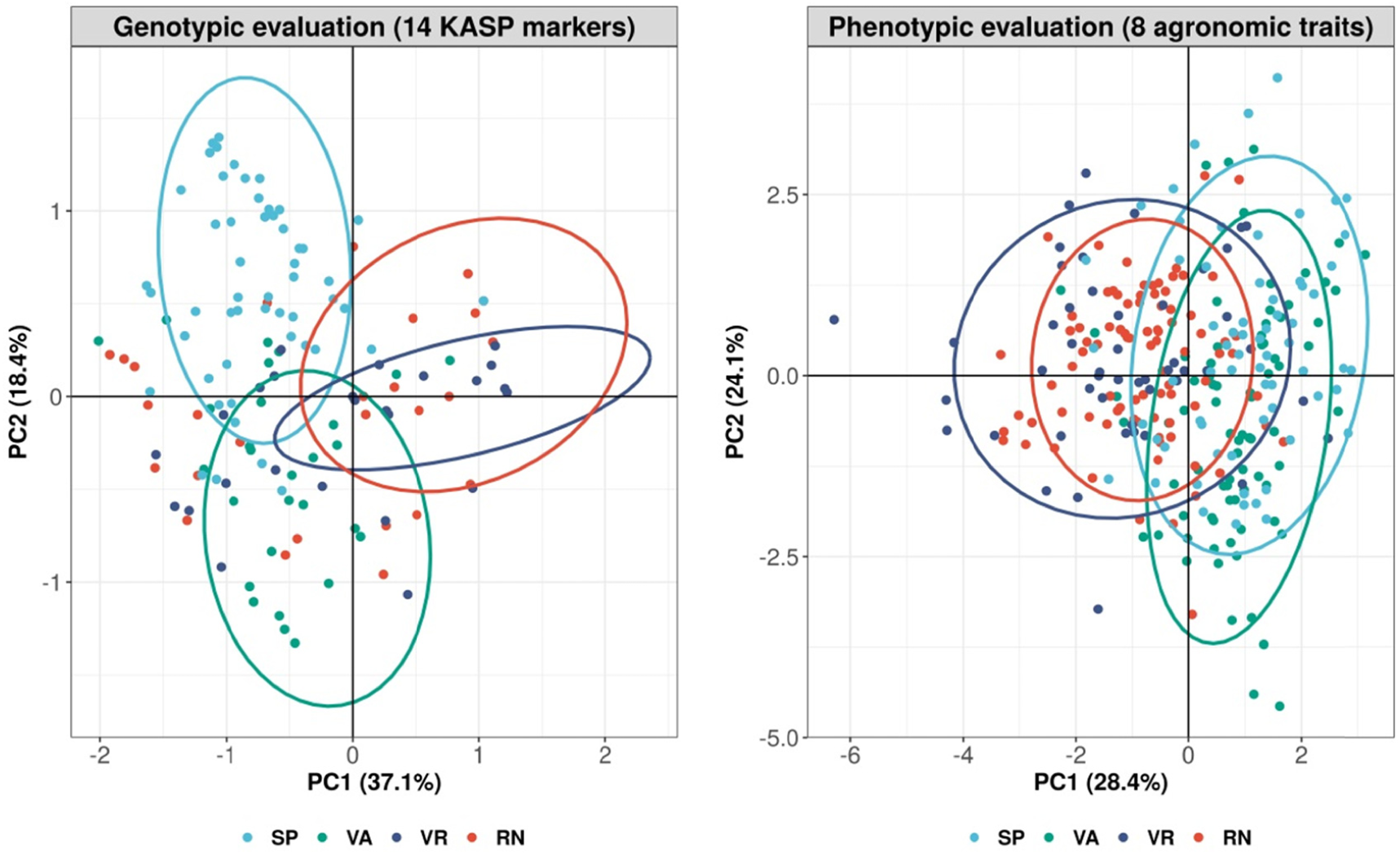
Principal component analysis (PCA) of 282 accessions based on genotyping data of 14 ko petitive allele-specific PCR (KASP) and phenotypic data. 282 accessions were visualized by 4 market types, Spanish (SP), Val**e**ncia (VA), Virginia (VR), Runner (RN).

**Figure 5.**
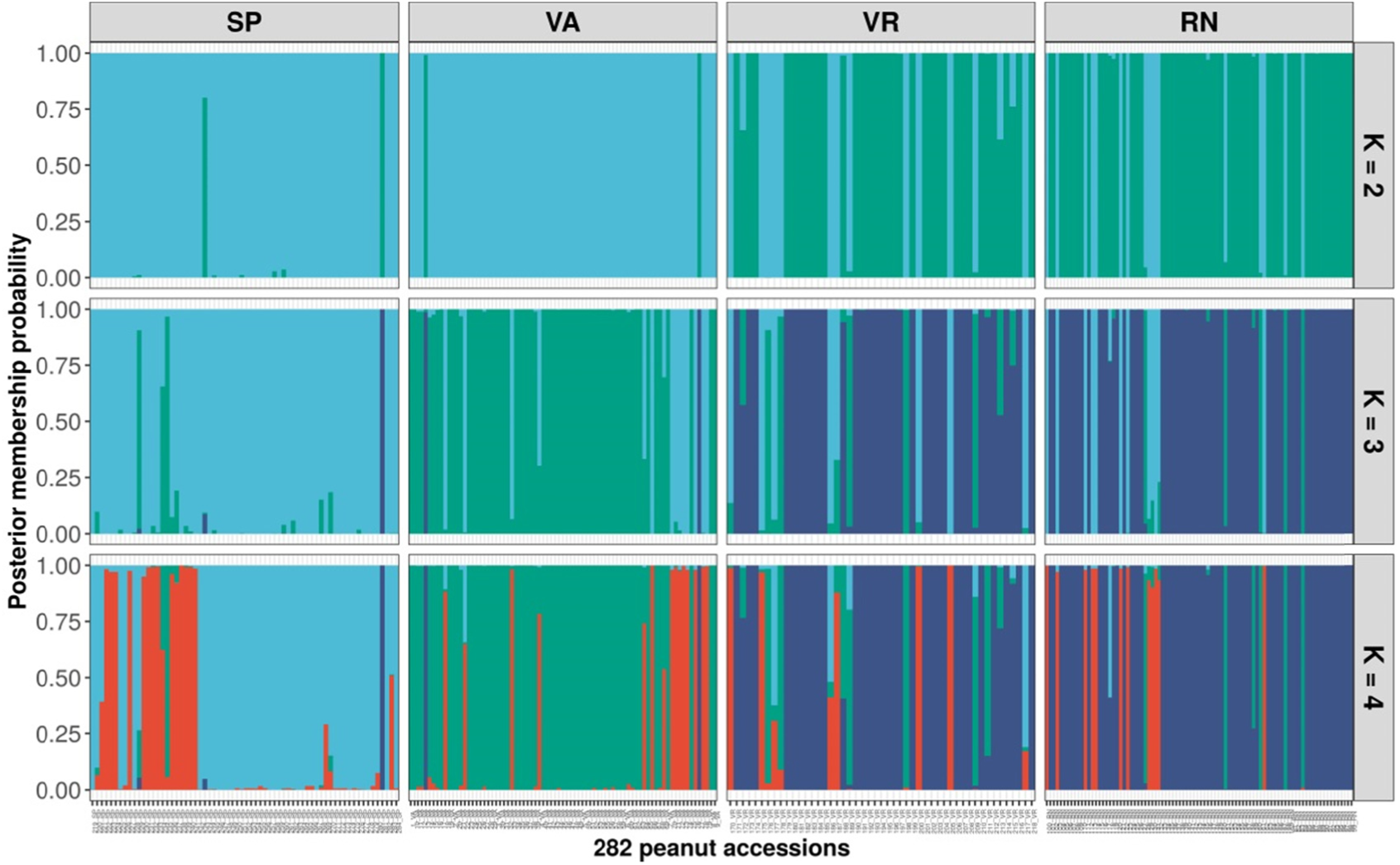
Bar plots of posterior membership probability produced by discriminant analysis of principal components (DAPC) for a number of clusters ranging from 2 to 4. The analysis was performed using 282 peanut accessions genotyped by 14 kompetitive allele-specific PCR (KASP) markers, visualized under the classification into the 4 market types: Spanish (SP), Valencia (VA), Virginia (VR), Runner (RN). The color code is the same as in Figure 4.

## Discussion

Molecular markers are of importance in many aspects of plant genetics and breeding, including variety identification, positional cloning, and the exploration of genetic diversity and population structure within germplasm. Developing molecular markers in the cultivated peanut was challenging because of its narrow genetic diversity and the sequences of high similarity between its two diploid genomes^38^. The assembled genome of cultivated peanuts and their diploid ancestors using NGS approaches has significantly boosted the genomic research in the peanut community. In particular, reduced-representation sequencing, such as GBS and RAD-seq, has been widely used in peanut research^20,21,22^. In this study, the RAD-seq approach was utilized to sequence 31 accessions of Taiwanese germplasm, that can be decomposed into a “global” subset containing 17 “introduced” accessions and a “local” subset containing 14 Taiwanese accessions, 12 being current elite cultivars, 1 being a landrace, and 1 being an advanced breeding line. The global subset has a higher average number of SNPs than the local subset, suggesting that the germplasm from abroad have more polymorphisms than the local germplasm, and this perspective was supported by further genetic diversity analyses and by performing clustering based on 3,474 out of 17,610 SNPs which can differentiate these 31 accessions.

Based on 3,474 SNPs, the genetic diversity of these 31 accessions was investigated using several approaches. As a whole, this panel has an average PIC, expected He, MAF and genetic distance of 0.16, 0.19, 0.86, 0.17, respectively, which is concordant with previous research using SNP genotyping^39,40,41^. In terms of the marker polymorphism, the PIC value is an indicator of markers that can reveal polymorphisms. Markers are considered as highly informative (greater than 0.5), reasonably informative (0.25 to 0.5) and only slightly informative (smaller than 0.25) according to their PIC values^29^. The average PIC of 0.16 from these 31 accessions falls in this last class, while 32% of the identified SNPs correspond to the reasonably informative class. In studies of other germplasm collections, the mean PIC value ranged from 0.08 to 0.26 even though some of these collections had a mixture of four market-type accessions including the U.S. Peanut mini core collection^39,40,41^, implying that the germplasm panel of 31 accessions chosen in this study preserves a high proportion of overall genetic diversity in spite of a smaller sample size. Concerning the origin of our germplasm, the global subset (n = 17) has a higher average PIC value (0.15) than the local subset (0.13 with n = 14). Similarly, the average He and genetic distance of the global subset (He = 0.19, distance = 0.17) are greater than that of local subset (He = 0.16, distance = 0.14), indicating that the global subset has larger genetic diversity than the local subset. While these 31 accessions were separated into three botanical varieties, accessions from subsp. *fastigiata* var. *fastigiata* (Valencia type) have larger He (0.18), PIC (0.14) and genetic distance (0.15) on average than subsp. *fastigiata* var. *vulgaris* (Spanish type, He = 0.13, PIC = 0.11, distance = 0.11) and subsp. *hypogaea* var. *hypogaea* (Virginia/Runner types, He = 0.12, PIC = 0.10, distance = 0.12). In the 31 accessions, 10 genotypes are from subsp. *fastigiata* var. *fastigiata* including 7 introduced accessions and 3 cultivars; in particular, 5 of 7 introduced accessions are from countries in South America including Brazil, Uruguay, Peru and Venezuela. The cultivated peanut originated from South America^42^, it is thus expected that genotypes of subsp. *fastigiata* var. *fastigiata* should have larger genetic diversity than genotypes of subsp. *fastigiata* var. *vulgaris* and subsp. *hypogaea* var. *hypogaea* composing most cultivars in Taiwan. They also are expected to have higher diversity than this study’s introduced accessions coming from regions away from the center of origin of cultivated peanut.

Genetic relationships among the 31 accessions were investigated using the pairwise genetic distance for the construction of the dendrogram and PCA (Supplementary Table S2). In general, these 31 accessions were grouped into three clusters in line with three botanical varieties. Group I and Group II with mainly subsp. *fastigiata* var. *fastigiata* and subsp. *fastigiata* var. *vulgaris*, respectively, were separated by a genetic distance of 0.19, and Group III was separated from Group I/II by a genetic distance of 0.21. These results indicate that Group I and Group II are more closely related to one-another than to Group III, in agreement with the botanical classification and with other studies having larger sample sizes^12,16,22^. This dendrogram result also suggests that the local cultivars in Taiwan might be suffering from low genetic diversity due to the excessive exploitation of narrow genetic resources as breeding material, notably genotypes of Spanish type germplasm. In the main clade within Group II, there are 9 accessions from the local subset containing 8 cultivars and 1 landrace. According to the pedigree of these 8 cultivars, many of them are genetically close to TNS9. TNS9 was developed in 1966 by pure line selection using the introduced line “Giay” from Vietnam, and this variety dominated more than 80% of peanut production in Taiwan in the 1980s because of its favorable flavor after roasting. This variety has also been widely exploited in peanut breeding programs in Taiwan, such as the development of TNG10, HL1, HL2, TN14 and TN18, all developed by the hybridization breeding method. Specifically, TNS9 was directly chosen as a parent of HL1, and indirectly contributed to the genetic background of the other four varieties by being selected as the parent of advanced breeding lines used in the breeding programs of these three varieties. Based on pedigree information, it is thus anticipated that HL2, a Valencia type variety, was grouped into the cluster with mostly Spanish type germplasm. On the other hand, TN16 and TN17, both rich in cyanidine-based anthocyanins on the seed coat, are 2 Taiwanese Valencia type cultivars derived from the same biparental breeding population using the hybridization of 2 landraces collected from central Taiwan. The dendrogram result showed that TN16 and TN17 were clustered into Group III; moreover, they are obviously separated from the other accessions in this clade. Therefore, these two varieties are not closely related to the three groups containing the other 29 accessions, suggesting that potential still exists to have locally collected genetic resources which can diversify the current Taiwanese germplasm. The same result of clustering was also discovered using PCA based on genetic distance (Fig. 2).

To understand the genetic diversity beyond the 31 accessions, at least within present conservation projects, and with in mind the improvement of peanut breeding programs in Taiwan, 14 sets of KASP markers were developed using the SNPs identified in our 31 accessions. We validated these KASP markers using 282 genotypes from the germplasm conservation center in TARI, and the PCA results showed that these genotypes were distinctly separated into 3 groups according to three botanical varieties (Fig. 4). This conclusion was also supported by K-means clustering, in particular with the choice k=3; beyond that value the BIC value does not improve much (Supplementary Fig. S2). In addition, these KASP markers clearly distinguished subsp. *fastigiata* and subsp. *hypogaea* accessions at k=2, and then separated var. *fastigiata* and var. *vulgaris* from the same subspecies *fastigiata* at k=3, which is compatible with previous work^22^. However, when setting k=4, the additional group has a mixture of four market types of germplasm belonging to all three botanical varieties, suggesting that exchanges of genetic background among these accessions may have occurred. This result of a fourth cluster not corresponding to subspecies or market types was also reported in other works^12,39^, and it may result from phenotyping difficulties^43,44^. In both sets, containing respectively 31 and 282 accessions, PCA was used to compare the effectiveness of molecular markers and phenotypic data to cluster samples, and it was demonstrated that PCA based on molecular markers provides more reproducible and satisfactory results than PCA based on phenotypic data (Fig. 2, Fig. 4, Supplementary Fig. S1).

Overall, the genetic diversity and relationship among germplasm in Taiwan was revealed by SNPs identified through the RAD-approach. Our analyses suggest that one should broaden genetic diversity by introducing novel germplasm to prevent genetic vulnerability. In addition, the KASP markers successfully developed here should be a useful tool for identifying the population structure of peanut germplasm collections or for conducting other genetic studies related or not to breeding.

## Supporting information

Supplementary File 1 (vcf format)

## Acknowledgements

We thank O.C. Martin, T. Blein and K.H. Chen for discussions and we are particularly grateful for the English editing from O.C. Martin. The authors also thank Dr. K.Y. Chen at National Taiwan University for providing resources and C.Y. Liu for the guidance while preparing RAD-seq libraries.

## Contributions

H.Y.D and S.S.W conceived the study and acquired the funding. Y.M.H designed the bioinformatics and data analysis pipeline and wrote the manuscript. S.R.L, H.F, W.C.H and H.I.K performed the experiments. Y.C.T supervised the student. All authors contributed to manuscript revisions.

## Data Availability

The sequencing data of 31 accessions produced in this study have been deposited at the NCBI BioProject PRJNA811600 (reviewer link https://dataview.ncbi.nlm.nih.gov/object/PRJNA811600?reviewer=9s5mq14otqrpnlq1f8amotftlu). All the codes related to this project are available in the github site https://github.com/ymhsu/ahdivertwn.

## References

1. Willett, W. et al. Food in the Anthropocene: The EAT-Lancet Commission on healthy diets from sustainable food systems. Lancet 393(10170), 447–492 (2019).

2. Barkley, N. A., Upadhyaya, H. D., Liao, B. & Holbrook C. C. Peanuts: Genetics, Processing, and Utilization. 67–109 (AOCS Press, 2016).

3. Desmae, H. et al. Genetics, genomics and breeding of groundnut (Arachis hypogaea L.). Plant Breeding 138(4), 425–444 (2019).

4. Moretzsohn, M. C. et al. Genetic diversity of peanut (Arachis hypogaea L.) and its wild relatives based on the analysis of hypervariable regions of the genome. BMC Plant Biol. 4(1), 11; https://doi.org/10.1186/1471-2229-4-11 (2004).

5. Oteng-Frimpong, R., Sriswathi, M., Ntare, B. R. & Dakora, F. D. Assessing the genetic diversity of 48 groundnut (Arachis hypogaea L.) genotypes in the Guinea savanna agro-ecology of Ghana, using microsatellite-based markers. Afr. J. Biotechnol. 14(32), 2484–2493. (2015).

6. Bertioli, D.J. et al. The genome sequences of Arachis duranensis and Arachis ipaensis, the diploid ancestors of cultivated peanut. Nat. Genet. 48, 438–446 (2016).

7. Zhuang, W. et al. The genome of cultivated peanut provides insight into legume karyotypes, polyploid evolution and crop domestication. Nat. Genet. 51(5), 865–876 (2019).

8. He, G. et al. Microsatellites as DNA markers in cultivated peanut (Arachis hypogaea L.). BMC Plant Biol. 3(1), 3; https://doi.org/10.1186/1471-2229-3-3 (2003).

9. Ferguson, M. E. et al. Microsatellite identification and characterization in peanut (A. hypogaea L.). Theor. Appl. Genet. 108(6), 1064–1070 (2004).

10. Mace, E. S., Phong, D. T., Upadhyaya, H. D., Chandra, S. & Crouch, J. H. SSR analysis of cultivated groundnut (Arachis hypogaea L.) germplasm resistant to rust and late leaf spot diseases. Euphytica 152(3), 317–330 (2006).

11. Tang, R. et al. Genetic Diversity in Cultivated Groundnut Based on SSR Markers. J. Genet. Genomics 34(5), 449–459 (2007).

12. Belamkar, V. et al. A first insight into population structure and linkage disequilibrium in the US peanut minicore collection. Genetica 139(4), 411–429 (2011).

13. Ren, X. et al. Genetic Diversity and Population Structure of the Major Peanut (Arachis hypogaea L.) Cultivars Grown in China by SSR Markers. PLoS ONE 9(2), e88091; https://doi.org/10.1371/journal.pone.0088091 (2014).

14. Bertioli, D.J. et al. The genome sequence of segmental allotetraploid peanut Arachis hypogaea. Nat. Genet. 51(5), 877–884 (2019).

15. Clevenger, J. et al. Genome-wide SNP Genotyping Resolves Signatures of Selection and Tetrasomic Recombination in Peanut. Mol. Plant. 10(2), 309–322 (2017).

16. Pandey, M. K. et al. Development and Evaluation of a High Density Genotyping ‘Axiom_Arachis’ Array with 58 K SNPs for Accelerating Genetics and Breeding in Groundnut. Sci. Rep. 7(1), 40577; https://doi.org/10.1038/srep40577 (2017).

17. Otyama, P. I. et al. Genotypic Characterization of the U.S. Peanut Core Collection. G3-Genes. Genom. Genet. 10(11), 4013–4026 (2020).

18. Baird, N. A. et al. Rapid SNP Discovery and Genetic Mapping Using Sequenced RAD Markers. PLoS ONE 3(10); e3376. https://doi.org/10.1371/journal.pone.0003376 (2008).

19. Elshire, R. J. et al. A Robust, Simple Genotyping-by-Sequencing (GBS) Approach for High Diversity Species. PLoS ONE 6(5), e19379; https://doi.org/10.1371/journal.pone.0019379 (2011).

20. Zhao, Y. et al. QTL mapping for bacterial wilt resistance in peanut (Arachis hypogaea L.). Mol. Breeding 36, 13; https://doi.org/10.1007/s11032-015-0432-0 (2016).

21. Han, S. et al. A SNP-Based Linkage Map Revealed QTLs for Resistance to Early and Late Leaf Spot Diseases in Peanut (Arachis hypogaea L.). Front. Plant Sci. 9, 1012; https://www.frontiersin.org/article/10.3389/fpls.2018.01012 (2018).

22. Zheng, Z. et al. Genetic Diversity, Population Structure, and Botanical Variety of 320 Global Peanut Accessions Revealed Through Tunable Genotyping-by-Sequencing. Sci. Rep. 8(1), 14500; https://doi.org/10.1038/s41598-018-32800-9 (2018).

23. Niu, C. et al. A safe inexpensive method to isolate high quality plant and fungal DNA in an open laboratory environment. Afr. J. Biotechnol. 7(16), (2008).

24. Subrahmanyam, P. et al. Screening methods and sources of resistance to rust and late leaf spot of groundnut. Information Bulletin 47, 24 (1995).

25. Etter, P. D., Bassham, S., Hohenlohe, P. A., Johnson, E. A. & Cresko W. A. SNP discovery and genotyping for evolutionary genetics using RAD sequencing. Methods Mol. Biol. 772, 157–178 (2011).

26. Catchen, J., Hohenlohe, P. A., Bassham, S., Amores, A. & Cresko W. A. Stacks: An analysis tool set for population genomics. Mol. Ecol. 22(11), 3124–3140 (2013).

27. Li, H. Aligning sequence reads, clone sequences and assembly contigs with BWA-MEM. Preprint at https://arxiv.org/abs/1303.3997v2 (2013).

28. Li, H. A statistical framework for SNP calling, mutation discovery, association mapping and population genetical parameter estimation from sequencing data. Bioinformatics 27(21), 2987–2993 (2011).

29. Botstein, D., White, R. L., Skolnick, M., & Davis, R.W. Construction of a genetic linkage map in man using restriction fragment length polymorphisms. Am. J. Hum. Genet. 32(3), 314–331 (1980).

30. Weir, B. S. Genetic data analysis II. (Sinauer Associates, 1996).

31. Lê, S., Josse, J., & Husson, F. FactoMineR: An R Package for Multivariate Analysis. J. Stat. Softw. 25(1), 1–18 (2008).

32. Prevosti, A., Ocaña, J. & Alonso, G. Distances between populations ofDrosophila subobscura, based on chromosome arrangement frequencies. Theor. Appl. Genet. 45(6), 231–241 (1975).

33. Kamvar, Z. N., Brooks, J. C. & Grünwald, N. J. Novel R tools for analysis of genome-wide population genetic data with emphasis on clonality. Front. Genet. 6, 208 https://doi.org/10.3389/fgene.2015.00208 (2015).

34. Jombart, T., Devillard, S. & Balloux, F. Discriminant analysis of principal components: a new method for the analysis of genetically structured populations. BMC Genet. 11, 94; https://doi.org/10.1186/1471-2156-11-94 (2010).

35. Jombart, T. & Ahmed, I. adegenet 1.3-1: New tools for the analysis of genome-wide SNP data. Bioinformatics 27(21), 3070–3071 (2011).

36. Paradis, E. & Schliep, K. ape 5.0: An environment for modern phylogenetics and evolutionary analyses in R. Bioinformatics 35(3), 526–528 (2019).

37. Wickham, H et al. “Welcome to the tidyverse.” J. Open Source Softw. 4(43), 1686; https://doi.org/10.21105/joss.01686 (2019).

38. Pandey, M. K. et al. Advances in Arachis genomics for peanut improvement. Biotechnol. Adv. 30(3), 639–651 (2012).

39. Otyama, P. I. et al. Evaluation of linkage disequilibrium, population structure, and genetic diversity in the U.S. peanut mini core collection. BMC Genomics 20(1), 481; https://doi.org/10.1186/s12864-019-5824-9 (2019).

40. Abady, S. et al. Assessment of the genetic diversity and population structure of groundnut germplasm collections using phenotypic traits and SNP markers: Implications for drought tolerance breeding. PLoS ONE 16(11), e0259883; https://doi.org/10.1371/journal.pone.0259883 (2021).

41. Nabi, R. B. S. et al. Genetic diversity analysis of Korean peanut germplasm using 48 K SNPs ‘Axiom_Arachis’ Array and its application for cultivar differentiation. Sci. Rep. 11(1), 16630; https://doi.org/10.1038/s41598-021-96074-4 (2021).

42. Weiss, E. A. Oilseed Crops. (Blackwell Science, 2000).

43. Holbrook, C. C. & Stalker, H. T. Plant Breeding Reviews. 297–356 (Wiley, 2002)

44. Barkley, N. A. et al. Genetic diversity of cultivated and wild-type peanuts evaluated with M13-tailed SSR markers and sequencing. Genet. Res. 89(2), 93–106 (2007).

