## Supplementary tables and figures for "Assessment of genetic diversity and SNP marker development within peanut germplasm in Taiwan by RAD-seq"

Supplementary Table S1. The characteristics of 31 accessions of the peanut germplasm used in this study

| Name | Botanical variety | Market type | Type | Origin | Important characteristics |
|---|---|---|---|---|---|
| PI153169 | var. *vulgaris* | Spanish | Introduced accession | Argentina | Pod rot resistance |
| PI259717 | var. *vulgaris* | Spanish | Introduced accession | Cuba | |
| PI565455 | var. *vulgaris* | Spanish | Introduced accession | United States | Early maturity |
| Tainung 7 (TNG7) | var. *vulgaris* | Spanish | Cultivar | Taiwan | High yield, large pod and sweet flavor |
| Tainung 10 (TNG10) | var. *vulgaris* | Spanish | Cultivar | Taiwan | High yield |
| Tainan 14 (TN14) | var. *vulgaris* | Spanish | Cultivar | Taiwan | High yield |
| Tainan 15 (TN15) | var. *vulgaris* | Spanish | Cultivar | Taiwan | High yield and large seeds |
| Tainan 18 (TN18) | var. *vulgaris* | Spanish | Cultivar | Taiwan | High yield |
| Tainan Selection 9 (TNS 9) | var. *vulgaris* | Spanish | Cultivar | Taiwan | High adaptability and yield |
| Hualieng 1 (HL1) | var. *vulgaris* | Spanish | Cultivar | Taiwan | High yield and tolerance to leaf chlorosis |
| India | var. *vulgaris* | Spanish | Introduced accession | India | |
| Xiamen | var. *vulgaris* | Spanish | Introduced accession | China | |
| Vietnam | var. *vulgaris* | Spanish | Introduced accession | Vietnam | |
| PI118480 | var. *fastigiata* | Valencia | Introduced accession | Brazil | Close to the center of origin |
| PI118989 | var. *fastigiata* | Valencia | Introduced accession | Brazil | Pod rot resistance |
| PI155112 | var. *fastigiata* | Valencia | Introduced accession | Uruguay | Pod rot resistance |
| PI314817 | var. *fastigiata* | Valencia | Introduced accession | Peru | Rust resistance |
| PI338337 | var. *fastigiata* | Valencia | Introduced accession | Venezuela | High yield |
| Tainan 16 (TN16) | var. *fastigiata* | Valencia | Cultivar | Taiwan | High anthocyanin in seed coats |
| Tainan 17 (TN17) | var. *fastigiata* | Valencia | Cultivar | Taiwan | High anthocyanin in seed coats |
| Hualieng 2 (HL2) | var. *fastigiata* | Valencia | Cultivar | Taiwan | High yield |
| E01001 | var. *fastigiata* | Valencia | Introduced accession | Japan | Large pod |
| E01004 | var. *fastigiata* | Valencia | Introduced accession | Mexico | |
| Red | var. *fastigiata* | Valencia | Landrace | Taiwan | Good aroma and large seeds |
| NS011001 | var. *hypogaea* | Virginia | Advanced breeding line | Taiwan | High yield |
| PI109839 | var. *hypogaea* | Virginia | Introduced accession | Venezuela | Early leaf spot resistance |
| Taichung 1 (TC1) | var. *hypogaea* | Virginia | Cultivar | Taiwan | High yield |
| PI145681 | var. *hypogaea* | Runner | Introduced accession | Egypt | Drought tolerance |
| PI599592 | var. *hypogaea* | Runner | Introduced accession | United States | High Oleic acid |
| PI203396 | var. *hypogaea* | Runner | Introduced accession | Brazil | Resistance of late leaf spot, southern blight and tomato spotted wilt virus |
| Penghu 1 (PH1) | var. *hypogaea* | Runner | Cultivar | Taiwan | Drought tolerance |

### Supplementary Table S2. The pairwise genetic distance between the 31 accessions

| | E01001 | E01004 | HL1 | HL2 | India | NS011001 | PH1 | PI109839 | PI118480 | PI118989 | PI145681 | PI153169 | PI155112 | PI203396 | PI259717 | PI314817 |
|---|---|---|---|---|---|---|---|---|---|---|---|---|---|---|---|---|
| E01004 | 0.047 | | | | | | | | | | | | | | | |
| HL1 | 0.200 | 0.199 | | | | | | | | | | | | | | |
| HL2 | 0.179 | 0.172 | 0.094 | | | | | | | | | | | | | |
| India | 0.198 | 0.196 | 0.114 | 0.105 | | | | | | | | | | | | |
| NS011001 | 0.151 | 0.147 | 0.147 | 0.122 | 0.149 | | | | | | | | | | | |
| PH1 | 0.227 | 0.223 | 0.190 | 0.198 | 0.221 | 0.208 | | | | | | | | | | |
| PI109839 | 0.233 | 0.227 | 0.187 | 0.199 | 0.226 | 0.210 | 0.067 | | | | | | | | | |
| PI118480 | 0.042 | 0.039 | 0.204 | 0.178 | 0.198 | 0.147 | 0.226 | 0.236 | | | | | | | | |
| PI118989 | 0.098 | 0.098 | 0.184 | 0.171 | 0.173 | 0.146 | 0.220 | 0.233 | 0.087 | | | | | | | |
| PI145681 | 0.227 | 0.223 | 0.188 | 0.200 | 0.221 | 0.209 | 0.074 | 0.028 | 0.234 | 0.232 | | | | | | |
| PI153169 | 0.205 | 0.200 | 0.120 | 0.118 | 0.095 | 0.157 | 0.220 | 0.229 | 0.202 | 0.177 | 0.228 | | | | | |
| PI155112 | 0.082 | 0.080 | 0.182 | 0.158 | 0.172 | 0.147 | 0.229 | 0.234 | 0.074 | 0.112 | 0.232 | 0.168 | | | | |
| PI203396 | 0.231 | 0.222 | 0.189 | 0.196 | 0.221 | 0.207 | 0.061 | 0.033 | 0.229 | 0.230 | 0.039 | 0.226 | 0.229 | | | |
| PI259717 | 0.199 | 0.194 | 0.115 | 0.117 | 0.083 | 0.152 | 0.215 | 0.219 | 0.197 | 0.172 | 0.220 | 0.083 | 0.169 | 0.219 | | |
| PI314817 | 0.057 | 0.059 | 0.204 | 0.175 | 0.200 | 0.148 | 0.227 | 0.237 | 0.053 | 0.096 | 0.235 | 0.197 | 0.089 | 0.234 | 0.196 | |
| PI338337 | 0.063 | 0.062 | 0.197 | 0.175 | 0.199 | 0.144 | 0.225 | 0.233 | 0.059 | 0.080 | 0.232 | 0.197 | 0.090 | 0.228 | 0.196 | 0.060 |
| PI565455 | 0.153 | 0.154 | 0.147 | 0.146 | 0.129 | 0.167 | 0.227 | 0.231 | 0.152 | 0.153 | 0.232 | 0.118 | 0.158 | 0.231 | 0.119 | 0.153 |
| PI599592 | 0.217 | 0.211 | 0.138 | 0.152 | 0.173 | 0.190 | 0.119 | 0.095 | 0.218 | 0.208 | 0.097 | 0.166 | 0.207 | 0.101 | 0.170 | 0.220 |
| Red | 0.199 | 0.199 | 0.039 | 0.097 | 0.116 | 0.147 | 0.189 | 0.190 | 0.202 | 0.185 | 0.187 | 0.122 | 0.178 | 0.185 | 0.117 | 0.206 |
| TC1 | 0.221 | 0.217 | 0.162 | 0.182 | 0.197 | 0.198 | 0.089 | 0.055 | 0.222 | 0.220 | 0.063 | 0.195 | 0.212 | 0.064 | 0.192 | 0.225 |
| TN14 | 0.195 | 0.193 | 0.044 | 0.097 | 0.121 | 0.151 | 0.185 | 0.186 | 0.198 | 0.183 | 0.182 | 0.122 | 0.178 | 0.182 | 0.121 | 0.199 |
| TN15 | 0.194 | 0.193 | 0.042 | 0.096 | 0.114 | 0.142 | 0.183 | 0.178 | 0.199 | 0.181 | 0.178 | 0.118 | 0.176 | 0.177 | 0.111 | 0.197 |
| TN16 | 0.244 | 0.242 | 0.210 | 0.211 | 0.237 | 0.223 | 0.170 | 0.169 | 0.246 | 0.244 | 0.171 | 0.237 | 0.243 | 0.165 | 0.227 | 0.247 |
| TN17 | 0.237 | 0.232 | 0.208 | 0.204 | 0.229 | 0.221 | 0.169 | 0.169 | 0.237 | 0.237 | 0.169 | 0.231 | 0.235 | 0.165 | 0.221 | 0.240 |
| TN18 | 0.187 | 0.180 | 0.095 | 0.104 | 0.109 | 0.139 | 0.182 | 0.180 | 0.188 | 0.175 | 0.178 | 0.133 | 0.170 | 0.177 | 0.121 | 0.190 |
| TNS9 | 0.210 | 0.207 | 0.119 | 0.111 | 0.091 | 0.131 | 0.211 | 0.215 | 0.212 | 0.188 | 0.212 | 0.117 | 0.189 | 0.211 | 0.108 | 0.210 |
| TNG10 | 0.200 | 0.192 | 0.060 | 0.097 | 0.111 | 0.148 | 0.176 | 0.175 | 0.200 | 0.179 | 0.173 | 0.130 | 0.177 | 0.173 | 0.124 | 0.202 |
| TNG7 | 0.203 | 0.200 | 0.091 | 0.119 | 0.125 | 0.158 | 0.177 | 0.171 | 0.204 | 0.186 | 0.173 | 0.124 | 0.186 | 0.173 | 0.127 | 0.201 |
| Vietnam | 0.201 | 0.195 | 0.121 | 0.102 | 0.058 | 0.147 | 0.221 | 0.223 | 0.196 | 0.174 | 0.220 | 0.105 | 0.176 | 0.218 | 0.091 | 0.199 |
| Xiamen | 0.208 | 0.201 | 0.116 | 0.115 | 0.099 | 0.154 | 0.206 | 0.206 | 0.206 | 0.183 | 0.202 | 0.115 | 0.184 | 0.202 | 0.093 | 0.207 |

Supplementary Table S2 (continued). The pairwise genetic distance between the 31 accessions

| | PI338337 | PI565455 | PI599592 | Red | TC1 | TN14 | TN15 | TN16 | TN17 | TN18 | TNS9 | TNG10 | TNG7 | Vietnam |
|---|---|---|---|---|---|---|---|---|---|---|---|---|---|---|
| E01004 | | | | | | | | | | | | | | |
| HL1 | | | | | | | | | | | | | | |
| HL2 | | | | | | | | | | | | | | |
| India | | | | | | | | | | | | | | |
| NS011001 | | | | | | | | | | | | | | |
| PH1 | | | | | | | | | | | | | | |
| PI109839 | | | | | | | | | | | | | | |
| PI118480 | | | | | | | | | | | | | | |
| PI118989 | | | | | | | | | | | | | | |
| PI145681 | | | | | | | | | | | | | | |
| PI153169 | | | | | | | | | | | | | | |
| PI155112 | | | | | | | | | | | | | | |
| PI203396 | | | | | | | | | | | | | | |
| PI259717 | | | | | | | | | | | | | | |
| PI314817 | | | | | | | | | | | | | | |
| PI338337 | | | | | | | | | | | | | | |
| PI565455 | 0.156 | | | | | | | | | | | | | |
| PI599592 | 0.215 | 0.186 | | | | | | | | | | | | |
| Red | 0.197 | 0.150 | 0.136 | | | | | | | | | | | |
| TC1 | 0.219 | 0.197 | 0.089 | 0.164 | | | | | | | | | | |
| TN14 | 0.196 | 0.151 | 0.135 | 0.033 | 0.164 | | | | | | | | | |
| TN15 | 0.193 | 0.145 | 0.134 | 0.034 | 0.153 | 0.042 | | | | | | | | |
| TN16 | 0.242 | 0.245 | 0.191 | 0.204 | 0.174 | 0.207 | 0.203 | | | | | | | |
| TN17 | 0.237 | 0.238 | 0.190 | 0.200 | 0.174 | 0.202 | 0.197 | 0.033 | | | | | | |
| TN18 | 0.188 | 0.151 | 0.145 | 0.100 | 0.165 | 0.100 | 0.096 | 0.204 | 0.197 | | | | | |
| TNS9 | 0.205 | 0.147 | 0.173 | 0.122 | 0.189 | 0.129 | 0.122 | 0.224 | 0.217 | 0.088 | | | | |
| TNG10 | 0.197 | 0.153 | 0.125 | 0.057 | 0.156 | 0.059 | 0.062 | 0.201 | 0.196 | 0.083 | 0.108 | | | |
| TNG7 | 0.199 | 0.147 | 0.142 | 0.089 | 0.151 | 0.092 | 0.087 | 0.188 | 0.184 | 0.123 | 0.138 | 0.102 | | |
| Vietnam | 0.195 | 0.136 | 0.173 | 0.124 | 0.203 | 0.128 | 0.124 | 0.232 | 0.225 | 0.110 | 0.094 | 0.119 | 0.131 | |
| Xiamen | 0.206 | 0.140 | 0.168 | 0.117 | 0.177 | 0.122 | 0.114 | 0.231 | 0.226 | 0.117 | 0.110 | 0.123 | 0.132 | 0.10 |

Supplementary Table S3. The primers of kompetitive allele-specific PCR (KASP) markers used in this study

| Marker ID | Primer_Allele_FAM | Primer_Allele_HEX | Primer_common |
|---|---|---|---|
| AHK_01 | GCAGTTTATTCAAGTTCTCACACAGC | GCAGTTTATTCAAGTTCTCACACAGA | GTGCTGCTAAACTGTCGGAAAGCAA |
| AHK_02 | CGGTTCTTGTATTGGATGAGGTT | CTCGGTTCTTGTATTGGATGAGGTA | CAGCTGCAGCCACCATGCTGAA |
| AHK_03 | GAGCTGCAGCAGCCACAAGAG | GAGCTGCAGCAGCCACAAGAA | CATGCCTGTGCCAAAGAAACGACAT |
| AHK_04 | TCATGTGGTTGAGCCTTCCAATG | GTTCATGTGGTTGAGCCTTCCAATA | GTGAATTCCTTGCTGAAGCGAGCAT |
| AHK_05 | AGAGTTGGTTAGTGAAGAAATGGTGA | GAGTTGGTTAGTGAAGAAATGGTGG | CTCCCATGTATCGTCAGTGATATTTGTAA |
| AHK_06 | AAACGGAAAGGGATGAGAAAACAGC | AAACGGAAAGGGATGAGAAAACAGG | GAGTTGCATGGAAAATTTCAAAGCTCAAAA |
| AHK_07 | CTTAGAAATCTCTCCTCTCCTCTC | CTCTTAGAAATCTCTCCTCTCCTCTA | CCCAAAGAGGCCCGTAAGAAACAAT |
| AHK_08 | GGAAGAGCTGAGAGGTTTCCAC | CGGAAGAGCTGAGAGGTTTCCAT | GTATGCAGAATGTCATGAACGCTTTCAAT |
| AHK_09 | GCGCATTATTGTTATTATTGTTCGGCT | CGCATTATTGTTATTATTGTTCGGCG | GGCATCGCTTCATCAGGATCTTCAT |
| AHK_10 | TCTTCTGTTTTGAATAACTATGTAGTTCG | ATTTCTTCTGTTTTGAATAACTATGTAGTTCA | ACATGTGTAGTTGCCATCTACACACAAAT |
| AHK_11 | ACTCTGCAGATGAACATGCATTTGTTT | CTCTGCAGATGAACATGCATTTGTTG | CATTGTCTTCAAAATAGGTCCTCTCTCTA |
| AHK_12 | TCATCACATTCGCCCATCAACTC | CTTCATCACATTCGCCCATCAACTT | GAGGAGGCACTCTGCTGCGTT |
| AHK_13 | CCTTCAACTATTTTCCATTTCGATCAC | ATCCTTCAACTATTTTCCATTTCGATCAT | TGCGGGATCGATAAATGCATGGGAA |
| AHK_14 | GTGTATTCTGATTCCCTTCTATACCAT | GTATTCTGATTCCCTTCTATACCAG | TCCAGGGAATGCAGAATCAATTGAGAAAT |

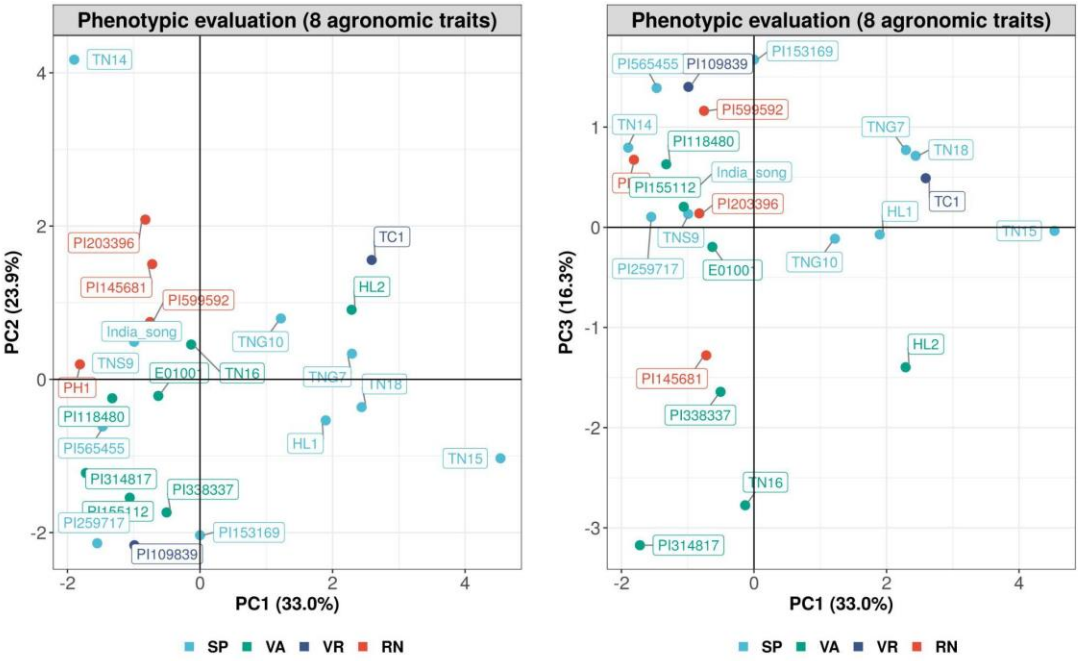


Supplementary Figure 1. Principal component analysis (PCA) of 24 accessions based on phenotypic data. 24 accessions were visualized by 4 market types, Spanish (SP), Valencia (VA), Virginia (VR), Runner (RN).

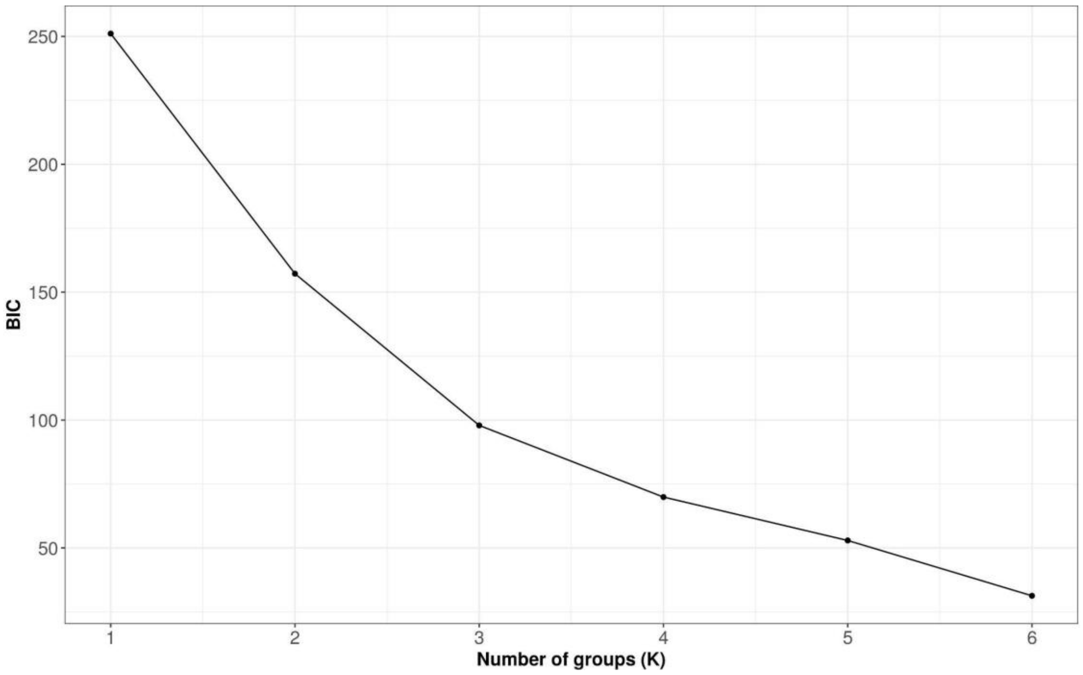

Supplementary Figure 2. The Bayesian information criterion (BIC) value with increasing number of groups (k) based on 282 accessions genotyped by 14 kompetitive allele-specific PCR (KASP) markers.
